# A Silk-Expressed Pectin Methylesterase Confers Cross-Incompatibility Between Wild and Domesticated Strains of *Zea mays*

**DOI:** 10.1101/529032

**Authors:** Yongxian Lu, Samuel A. Hokin, Jerry L. Kermicle, Mathew M. S. Evans

## Abstract

Despite being members of the same species, some strains of wild teosinte maintain themselves as a distinct breeding population by blocking fertilization by pollen from neighboring maize plants. These teosinte strains may be in the process of evolving into a separate species, since reproductive barriers that block gene flow are critical components in speciation. This trait is conferred by the *Teosinte crossing barrier1-s (Tcb1-s)* haplotype, making *Tcb1* a speciation gene candidate. *Tcb1-s* contains a female gene that blocks non-self-type pollen and a male function that enables self-type pollen to overcome that block. The *Tcb1-female* gene encodes a Pectin Methylesterase, implying that modification of the pollen cell wall by the pistil is a key mechanism by which these teosinte females reject foreign (but closely related) pollen.

**One sentence summary:** The *Tcb1-female* gene encodes a Pectin Methylesterase that in teosinte silks prevents fertilization by maize pollen.

## Main text

Maize (*Zea mays* ssp *mays*) was domesticated from annual teosinte (*Zea mays* ssp *parviglumis*) in the Balsas River valley of Mexico (*1*). In some locations, sympatric populations of domesticated maize and annual teosinte grow in intimate associate and flower synchronously, but rarely produce hybrids (*2, 3*). In sexually reproducing plants, reproductive barriers exist at different stages, including pre-pollination, post-pollination, and post-fertilization. Post-pollination barriers depend on interaction between the pollen grain and the female reproductive organs (stigma, style, and ovule). In *Zea mays*, haplotypes at three loci, *Gametophyte factor1-s (Ga1-s), Gametophyte factor2-s (Ga2-s)*, and *Teosinte crossing barrier1-s (Tcb1-s)*, confer Unilateral Cross-Incompatibility. While *Ga1-s* and *Ga2-s* are widespread in domesticated maize, *Tcb1-s* is almost exclusively found in wild teosinte populations. The *Tcb1-s* haplotype, like *Ga1-s* and *Ga2-s*, confers unilateral cross-incompatibility against varieties carrying the *tcb1* (or *ga1* or *ga2*) haplotype. Viewed otherwise, *Tcb1-s* provides a pollen function that overcomes the crossing barrier. The latter view is preferred since pollen containing both *Tcb1-s* and *tcb1* haplotypes fertilizes *Tcb1-s* plants, indicating that *Tcb1-s* compatibility is not overcome by the *Tcb1-s* : *tcb1* mismatch, as is also the case for the *Ga1* and *Ga2* systems (*4, 5*). *Tcb1-s* was first described in teosinte subspecies *mexicana* Collection 48703 from the central and southern Mexico; this strain also contained the male-only haplotype, *Ga1-m*, of the *Ga1* locus which together with male and female functions of *Tcb1*, make up the Teosinte Incompatibility Complex (TIC) (*2, 3*).

Collections of teosinte of both *mexicana* and *parviglumis* subspecies from the central Mexican plateau carry *Tcb1-s (*6*). Tcb1-s* confers to females the ability to block fertilization by maize (*tcb1* type) pollen by restricting pollen tube growth (*7*). In the reciprocal cross, teosinte pollen is able to fertilize maize, although poorly when in competition with maize pollen (*3*). *Tcb1* was proposed to be a candidate speciation gene contributing to isolation of diverging maize and teosinte populations, as wild teosinte populations respond to the pressure of cultivated, closely related varieties of domesticated maize (*6*).

The male and female functions of *Tcb1-s* are tightly linked but separable by recombination (*7*). Thus, there are four functional classes at this locus (Table S1 for gene content and origin): *Tcb1-s* has both functional male and female genes, *Tcb1-m* has only the functional male gene (*6, 7*), *Tcb1-f* has only the functional female gene, and the *tcb1* haplotype found in almost all maize lines has neither of the two functional genes. In teosinte, *Tcb1-f* activity in the silks prevents fertilization by maize *(tcb1)* pollen, while *Tcb1-m* activity in pollen enables fertilization of *Tcb1-f* females (*7*).

To clone the *Tcb1* genes, fine mapping of *Tcb1-s::Col48703* haplotype was performed based on a *tcb1* backcross population with a population of approximately 15,000 chromosomes. Using maize B73 genome as a reference (*8*), the *Tcb1* locus was delimited to a region spanning 480 kb on the short arm of chromosome 4. Within this region, there are eleven annotated genes. However, all of these were ruled out as candidates for *Tcb1* functions because they either had identical sequence with identical expression levels between *tcb1* and *Tcb1-s* haplotypes or no expression in the silk or pollen in *Tcb1-s* or *tcb1* (mapping markers included in Table S2). The *Tcb1* genes, therefore, are likely absent from the maize genome. This is not surprising considering the widespread structural variations in genomes between maize lines and between teosinte populations (*9*).

To identify *Tcb1-f* knockout mutants, maize lines homozygous for the *Tcb1-s::Col48703* haplotype and carrying active *Mutator* transposons were crossed to maize inbred A195 *su1*. The progeny are expected to be heterozygous for *Tcb1-s* with *su1* approximately 6 cM away and in repulsion (*3*). Due to the rejection of the *tcb1* pollen (which is predominantly su1), about 3% of the kernels in every ear with functional *Tcb1-f were* expected to be *sugary* in this open-pollinated population, while any ears without a crossing barrier were predicted to segregate *su1* at 25%. Out of a population of approximately 6,000 individuals, two exceptional ears were found. One ear segregated for 25.6% sugary. This allele is termed *tcb1-f(KO1)*. The second isolate contained a sector of about 45 kernels within which the segregation was one-fourth sugary despite sugary segregating at ~3% over the rest of the ear. This allele is termed *tcb1-f(KO2)*. Mixed pollination tests with the progeny of both individuals show that the loss of function is heritable, and both variants fertilized a *Tcb1-s/tcb1* strain normally, indicating the retention of the male function of *Tcb1-s (Tcb1-f* mutated, but *Tcb1-m* intact) (Fig. 1). In the case of *tcb1-f(KO2)*, progeny of seeds within the loss-of-function side of the ear inherited the knock-out, while those on the other side of the ear inherited fully functional *Tcb1-s*.

**Fig. 1.**
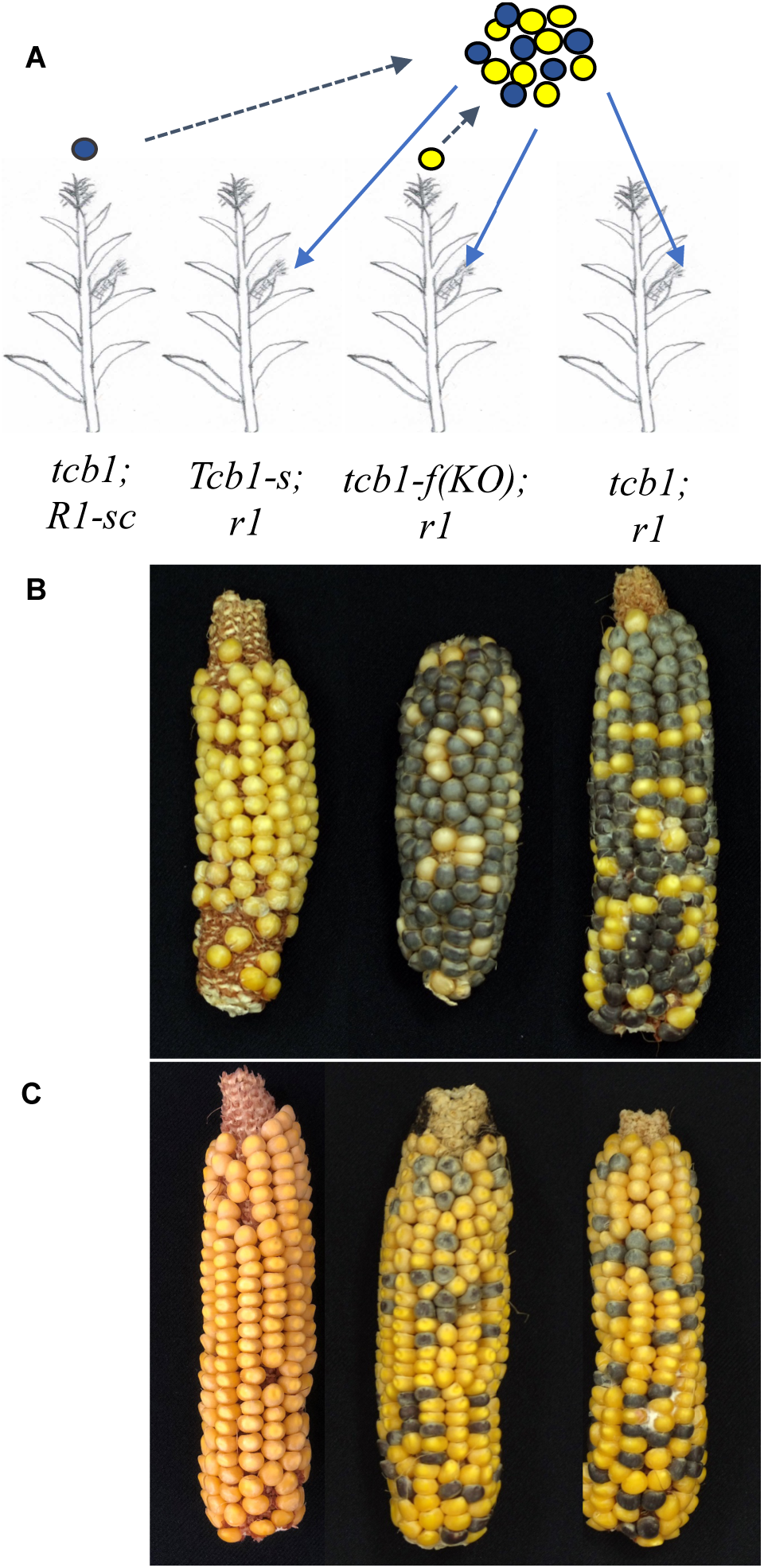
Mixed pollination test of the *Tcb1-f* mutants. (A) scheme of the experiment: Two pollen donor lines and three pollen receiver lines were used. Pollen from a *tcb1; R1-self color* maize line (purple circles), produces purple kernels, while pollen from test plants (yellow circles) produces white or yellow kernels. Pollen from the two donors was mixed and put on three receiving ears: (*1*) *Tcb1-s* ears to verify the *Tcb1-male* function from the *KO* line; (*2*) *tcb1-f(KO)* ears to test the presence/absence of the female barrier in the knockout mutant individuals; and (*3*) a maize *(tcb1; r1)* neutral ear to identify the ratio of viable pollen grains from the two donors in the mixture. (B) Ears from the three pollen recipients for the *tcb1-f(KO1))* test. (C) Ears from the three pollen recipients for the *tcb1-f(KO2)* test. In both tests, pollen from the *KO* plants successfully fertilized the *Tcb1-s* ear (left ear in B and C), while the ears from *KO* plants showed no barrier to *tcb1* maize pollen with a similar frequency of purple kernels on mutant ears and the neutral test ears (middle and right ears in B and C).

RNA from silks of four genotypes were subjected to short read RNA-seq. Transcript models were assembled *de novo* from the RNA-seq reads, and expression levels of genes were compared between these two knockout mutants, a standard maize inbred line W22 (genotype *tcb1*), and a functional *Tcb1-s* line (a W22 subline to which the *Tcb1-s::Col48703* haplotype had been introduced by backcrossing). One gene, named here *Pertunda* (Roman fertility goddess who enables penetration, parallel to the control of pollen tube penetration in pistils), encoding a maize Pectin Methylesterase38 (PME38) homolog, was identified as a candidate for the *Tcb-f* gene. *Pertunda* is highly expressed in *Tcb1-s* silks (with a peak read depth of ~100,000) compared to the standard maize *tcb1* W22 silks, *tcb1-f(KO2)* silks (maximum read depths of ~100) and *tcb1-f(KO1)* silks (maximum read depth of ~10,000 for the 5’ end and ~100 for the 3’ end of the transcript model) (Fig.2a). Based on the mRNA sequence, PCR primers were designed to isolate a BAC (Bacteria Artificial Chromosome) clone from a library we constructed from a maize line to which the *Tcb1-s::Col48703* haplotype had been introduced by backcrossing. By comparing mRNA and gene sequences, a 99-base intron was identified in *Pertunda*, which explains the gap between the two signal peaks in *Tcb1-s*. The intron region showed the same level of expression as that from the whole gene region in *tcb1-f(KO2)and* in the W22 maize line and in the downstream gene region in *tcb1-f(KO1)*. qRT-PCR confirmed this expression difference (Fig.2b). *Pertunda* is not present in the maize B73 reference genome, which is consistent with the mapping data, and the closest homologs of *Pertunda* are located at the *ga1* locus (*10*).

**Fig. 2.**
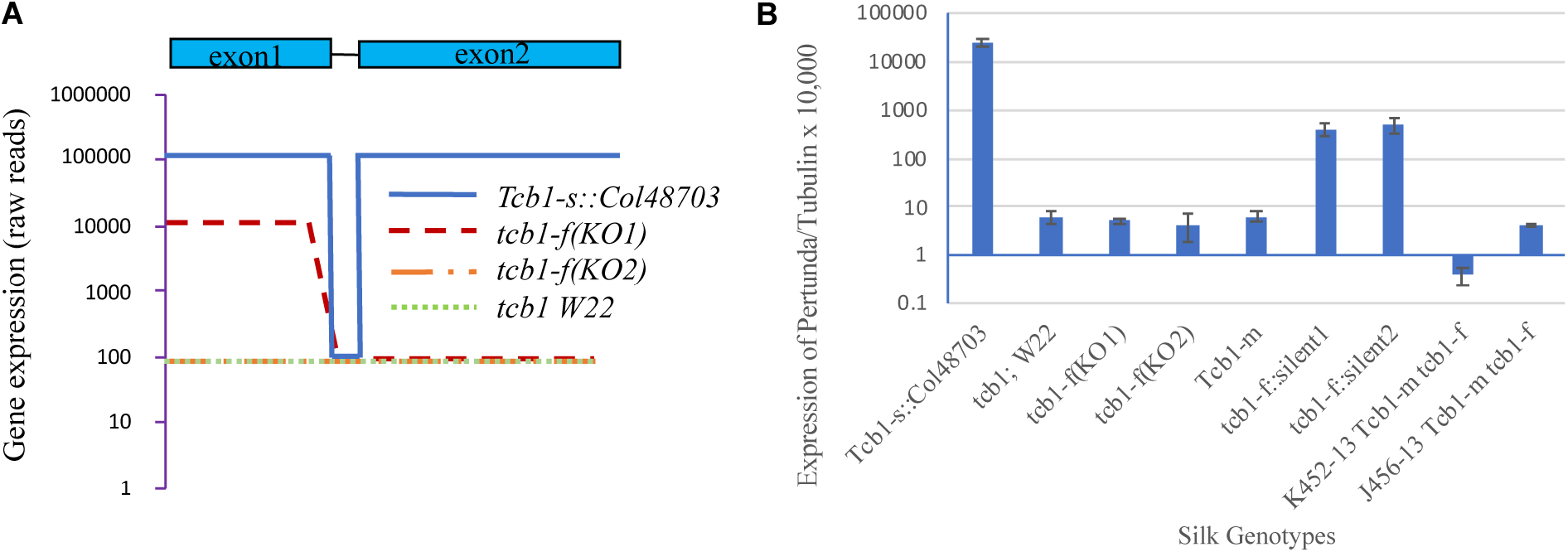
Gene expression profiling identifies *Pertunda* as a *Tcb1-f* candidate gene. RNA samples collected from silks of different genotypes was analyzed by RNA-Seq and RT-PCR. (A) *Pertunda* gene structure is shown above the graph (solid line indicates single intron), and RNA-Seq read depth is shown for the *Pertunda* gene. (B) *Pertunda* gene expression, compared to *tubulin* levels as a control, as measured by qRT-PCR in a variety of loss-of-function *Tcb1-s* lines: *Tcb1-s*, full strength *Tcb1-s* barrier line; *tcb1-f(KO1)* and *tcb1-f(KO2)*, two loss-of-function alleles from a *Mutator* transposon mutagenesis; *Tcb1-m*, a spontaneous *Tcb1-male* only line; *tcb1-f::silent lineage1* and *tcb1-f::silent lineage2*; and *K452-13* and *J456-13*, two *Tcb1-male* only lines that lost *Tcb1-f* by recombination.

In addition to the two knockout mutants from the active *Mutator* transposon population, several additional lines derived from the *Tcb1-s::Col48703* accession have lost female barrier function. One was recovered during early backcrossing of the *Tcb1-s::Col48703* haplotype into maize (*2*). Mixed pollination confirmed this is a *Tcb1-male* only plant (Fig.S1). Additionally, two independent *Tcb1-s* lines were isolated in which the barrier gradually lost the strength during ten generations of backcrossing to maize and selection for *Tcb1-male* function (*7*). These two lines were named as *tcb1-f::silent lineage1 (tcb1-f::sl1)* and *tcb1-f::silent lineage2 (tcb1-f::sl2)*, based on the “progressive” manner of the barrier weakening. *Pertunda* has much lower expression in these three additional *tcb1* lines, as shown with qRT-PCR (Fig. 2 b). Expression of *Pertunda* was also tested on the two *Tcb1-m* recombinants from the mapping population, which have lost the *Tcb1-female* gene by recombination. Again, *Pertunda* expression was much lower than in *Tcb1-s* lines (Fig. 2 b).

Using a PCR-based dCAPS (Derived Cleaved Amplified Polymorphic Sequence) marker designed for the *Pertunda* gene, it was shown that *Pertunda* maps to the *tcb1* locus (Fig. S2). This marker was then tested on the fifteen closest recombinants from the mapping population of ~15,000 individuals (including four recombinants between the *Tcb1-f* and *Tcb1-m* genes) (*7*). Of the fifteen plants, six carried *Tcb1-f* and blocked maize pollen, and nine lacked the barrier. Results showed that all the six recombinants that carry the barrier had the *Pertunda* gene, while in all nine recombinants that are receptive to maize pollen, *Pertunda* was absent. This shows a perfect physical linkage between *Pertunda* and the *Tcb1-f* barrier function. RNA-seq data suggest that the mutation in *Pertunda* occurred somewhere in the first exon in the *tcb1-f(KO1)* mutant (Fig. 2a). PCR data confirmed there was a disruption within the coding region of *Pertunda in KO1* (Fig. S3). Quite differently, in *tcb1-f(KO2)* mutant silk RNA-seq reads had the same low level of expression as *tcb1* silks along the whole *Pertunda* transcript. Whole genome resequencing of both mutants identified a *Hopscotch* retrotransposon insertion in the first exon in *tcb1-f(KO1)*, close to the site where *Pertunda* expression drops sharply. PCR spanning both ends of the insertion confirmed the insertion event and the border sequences (Fig. S4). In contrast, in *tcb1-f(KO2), Pertunda* was fully assembled, consistent with the PCR data that the coding region is present (Fig. S3). The *tcb1-f(KO2)* allele then could either be mutated in a regulatory region, potentially hundred kilobases away from the coding region, or could be an epi-allele. Similarly, no mutations were found in the coding region of *Pertunda* in the *Tcb1-m* line or the *tcb1-f::sl1* or *tcb1-f::sl2* lines described above.

The *tcb1-f(KO2), tcb1-f::sl1*, and *tcb1-f::sl2* lines were tested for reversion to *Tcb1-s* in double mutants with *mediator of paramutation1 (mop1)* mutation. *MOP1* encodes a RNA-dependent RNA polymerase and is a key component of RNA-directed DNA Methylation (*11*). *mop1* mutations reactivate silenced genes and affect broad developmental programs (*12*). Re-activation of the *Tcb1-f* function was rare; in only ~14-22% of the *mop1* females tested, did the loss-of-function plants show some recovery of *Tcb1-f* function. Pollen competition experiments were performed for full strength *Tcb1-s* females, *tcb1* females, and the *tcb1-f* loss of function lines without sequence changes (primarily *tcb1-f(KO2), tcb1-f::sl1*, and *tcb1-f::sl2)* (Fig. 3). All of the *Tcb1-s* ears tested showed strong preference for *Tcb1-s* pollen (0-7% kernels from *tcb1* pollen regardless of the ratio of the two pollen types in the mix as indicated by the neutral ear) with the kernel ratio on the test ear and control ear being different from each other at p<0.0001 (Fisher exact test) (Fig 3b). Of the 36 *mop1; tcb1-f loss of function* females tested only one had as strong of a pollen preference as full strength *Tcb1-s* females, but five had a difference between the test and control ears at p<0.0001 and an additional three females could be included if the stringency was relaxed to p<0.01 (Fig. 3b). These partial revertants included plants of lines *tcb1-f(KO2), tcb1-f::sl1*, and *tcb1-f::sl2*. Of the twelve loss of function plants tested that were heterozygous wild-type for *mop1*, none of the plants passed the more stringent p<0.0001 threshold and one passed the less stringent p<0.01 threshold. It may be that maintaining the silenced derivatives of *tcb1-f* with *mop1* for multiple generations would increase the revertant frequency.

**Fig. 3.**
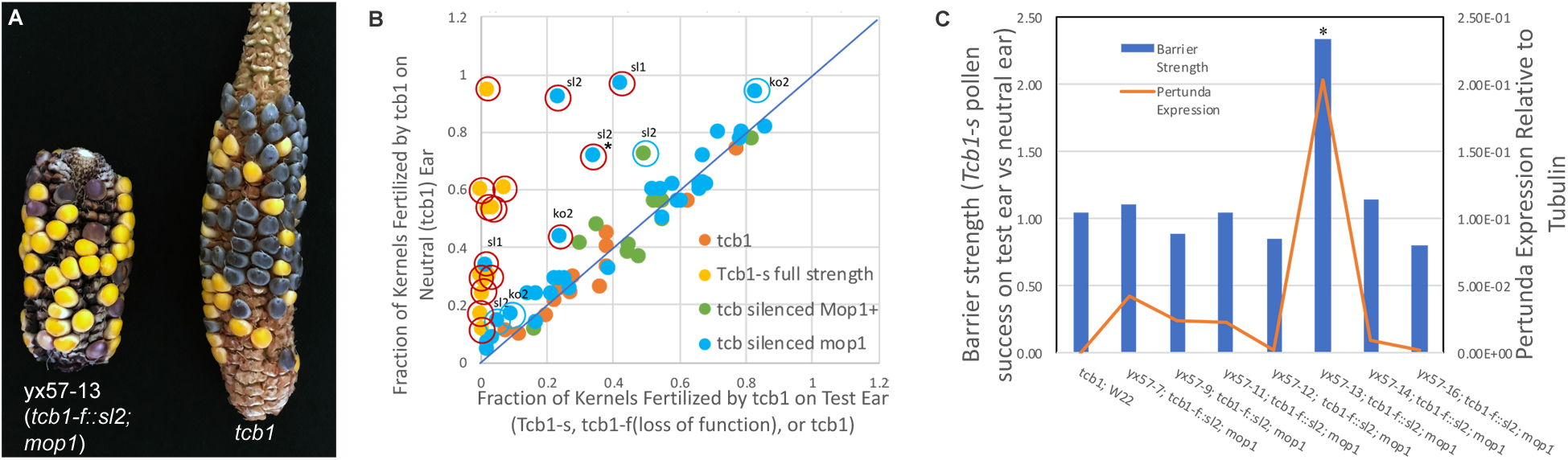
Reversion of *tcb1-f* loss-of-function. (A) Revertant ear (marked by asterisk in B and C) pollinated by a mix of *tcb1; R1-sc* and *Tcb1-m*; *r1* pollen showing higher frequency of yellow kernels on the test ear than the *tcb1* control ear. (B) Results of mixed pollination tests on four genotypes: *tcb1, Tcb1-s, tcb1-f(loss-of-function(lof))* alleles, and *tcb1-f(lof) mop1* double mutants. The percentage of kernels from *tcb1* pollen on the test ear are plotted on the x-axis and the percentage of kernels from *tcb1* pollen on the control *tcb1* ear on the y-axis. Equal percentages in the two ears indicates no barrier (line with slope=1). Red circles indicate ears with significantly fewer *Tcb1* kernels in the test ear vs. the control ear at p<0.0001 and blue circles at 0.0001<p<0.01. The loss-of-function line is indicated for each revertant. (C) *Pertunda* gene expression level and barrier strength. Barrier strength is expressed as the ratio of kernels from *tcb1-s* vs *tcb1* pollen on the test ear vs. the control *tcb1* ear (Columns), with a fraction of 1.0 indicating no barrier and significantly higher values a functional barrier. *Pertunda* RNA levels are expressed as relative to the *tubulin* control gene (Orange Line).

A subset of homozygous *mop1 tcb1-f::sl2* plants were tested at random for *Pertunda* expression in silks prior to pollination. Among the seven tested plants, one plant, yx57-13, showed about four hundred fold higher expression compared to that of the standard W22 maize and eight times higher than *tcb1-f::sl2* plants (Fig. 3C). This plant was the only one of those tested for *Pertunda* expression that recovered the ability to reject *tcb1* pollen, although not as efficiently as full strength *Tcb1-s* plants, which have still higher expression of *Pertunda* than this revertant. This indicates a correlation between *Pertunda* expression level and the female barrier strength, and further supports *Pertunda* as the *Tcb1-f* gene.

In addition to the *Tcb1-s::Col48703* strain descried above, three other teosinte-derived *Tcb1-s* lines, two from ssp. *mexicana* and one ssp. *parviglumis* (*6*), were tested for *Pertunda* expression in silk tissue. In all three lines, *Pertunda* expression levels are extremely high and comparable to that of the original central plateau *TIC* haplotype *Tcb1-s::Col48703* (Fig. 4). Interestingly, even though none of the modern north American maize lines tested to date carry the *Tcb1-s* haplotype, an ancient Maiz Dulce variety, Jalisco78 line 1222-2, grown at intermediate altitudes in southwestern Mexico carries *Tcb1-s* (*13*). This is a specialty line that may have undergone selection for cross-incompatibility factors similarly to *Ga1-s* in maize popcorn lines. Whether this Maiz Dulce line acquired *Tcb1-s* from nearby teosinte populations during its origin is unknown, but maize lines from this region have been shown to have substantial introgression from *mexicana* teosintes (*14*). Predicted *Pertunda* coding sequences are identical in all five *Tcb1-s* lines: three *mexicana* accessions, one *parviglumis* accession, and the Maiz Dulce line. One Single Nucleotide Polymorphism (SNP) in the intron separates these lines into two groups: one group including the *parviglumis* line (Col104-4a) and one *mexicana* line (Col109-4a), and the other group including two *mexicana* lines (Col48703 and Col207-5d) and the Maiz Dulce line (Fig. S5).

**Fig. 4.**
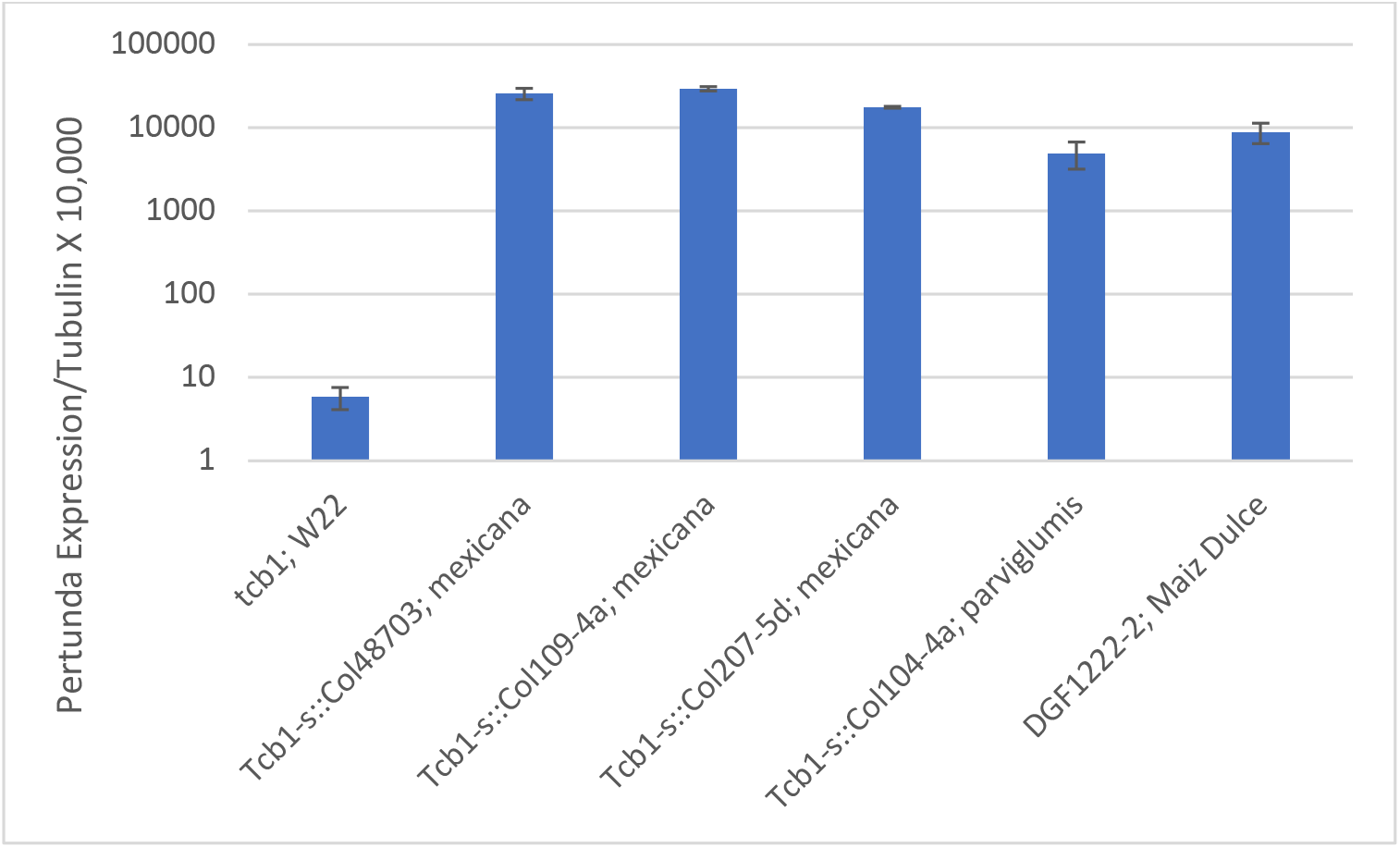
Expression of *Pertunda* in different *Tcb1-s* lines collected in different locations in Mexico (*6*). W22 is a standard maize *Tcb1* line, *Tcb1-s::Col48703, Tcb1-s::*109-4a, and *Tcb1-s::207-5d* are three independent collections of *Zea mays ssp. mexicana* tesointe lines; *Tcb1-s::*104-4a, a *Zea mays ssp. parviglumis* line; and DGF1222-2 from Maiz Dulce, an ancient Mexican maize sweet corn variety (*13*).

The most similar gene to *Pertunda* is a candidate PME gene for *Ga1-female* function. This gene, termed *ZmPME3*, was found to be expressed in the silks of *Ga1-s*, but not in *ga1* silks, and *ZmPME3* was located to the *Ga1* mapping region(*10*). Alignment of the *ZmPME3* and *Pertunda* show that the two PMEs differ in nine amino acids (Fig. S6). The number of polymorphisms (15 of 1296 nucleotides) between *Pertunda* and *ZmPME3* suggests that these two genes diverged approximately 175,000 years ago, well before the split between the *mexicana* and *parviglumis* subspecies of teosinte and just before the split between *Zea mays parviglumis* and *Zea luxurians*, using calculated nucleotide substitution rates for maize (*15*) and a calculated time since the split between *mexicana* and *parviglumis* of ~60,000 years and *parviglumis* and *luxurians* of ~140,000 years (*16*). It will be interesting to test whether the *Tcb1-male* and *Ga1-male* genes diverged at a similar time, suggesting they were already adjacent before divergence.

The *Tcb1* and *Ga1* barriers may share a similar mechanism, but because they are mostly cross-incompatible with one another they likely differ in their interacting partners. However, *Tcb1-s* and *Ga1-s* are not fully cross-incompatible. In situations where pollen rejection is not absolute, *Tcb1-s* pollen has a competitive advantage over *tcb1* pollen on *Ga1-s* or *Ga2-s* silks. This is true for all combinations of interactions between crossing barrier loci (*3, 5*) and is consistent with them encoding related proteins, although the behavior of pollen tubes during rejection by each system is slightly different (*7*). *Pertunda* encodes a group 1 type of PME without an N-terminal Pectin Methylesterase Inhibitor (PMEI) domain (*17*), and contains a predicted signal peptide, so it has the potential to be secreted and interact directly with the pollen tube to remove methyl-esters from the pectin wall of the pollen tube. Esterified pectins are typically associated with the tip of the growing pollen tube, while de-esterified pectins are enriched distally, and there is a correlation between pectin de-esterification and increased cell wall stiffness (*18*). Pollen tubes have a “soft tip-hard shell” structure, in that the tip region of the tube cell wall has a single pectin layer that is strong enough to withstand turgor pressure, but plastic enough to allow cell expansion and growth (*19*). Inside pollen tubes, pectin is synthesized and esterified in Golgi compartments before delivery to the tip cell wall via vesicle trafficking (*20*), where it can be de-esterified by PMEs (*21*). Pollen cells finely tune the stiffness of the tip cell wall to sustain pollen tube elongation. Either under-or over-supply of PME activity can result in disturbed pollen tube growth and compromised male fertility (*22–25*). The PERTUNDA (and ZMPME3) protein falls into the Plant 1a clade of mature PME enzymes (*26*) (Fig. S7).

In summary, genetic and genomic data identify *Pertunda* as the *Tcb1-female* barrier gene. Teosinte lines carrying *Pertunda* block maize pollen that lacks the male function provided by *Tcb1-m*. That the *Tcb1-f* gene encodes a cell wall modifying enzyme is consistent with the model that incompatibility with *tcb1* occurs via incongruity rather than active targeting of a *Tcb1-m* encoded protein (*4*). It will be interesting to test how universal this barrier mechanism is among sexual reproducing plants. Surprisingly, it was shown that another PME family member is encoded by the *Ga1-male* gene (*27*) (in a very distinct clade, Plant X2, of PME enzymes), raising the possibility that the biochemical barrier to pollen and the ability of pollen to overcome that barrier are conferred by different classes of PME proteins.

The grass family is known to have widely distributed self-incompatibility (SI) among species, however, the molecular nature of the SI genes and how it is related to interspecific cross-incompatibility are not known(*28, 29*). The grasses also have an unusually high species diversity for a family with abiotic pollinators (*30*). Identification of the *Tcb1-female* gene may facilitate research into the mechanisms of speciation in the grasses. Agriculturally, this work may help managing specialty crop populations by preventing pollen contamination. It may also facilitate development of breeding tools to enrich crop genetic pools by backcrossing crops to their ancestors for the purposes of yield increase or enhanced stress resistance.

## Acknowledgements

The authors would like to thank Kathy Barton for the support that kept this project alive, encouragement, and insight and members of the Evans and Barton labs past and present for helpful discussions. We would also like to thank Beverly Oashgar for help with the screen for loss-of-function mutants and David Heller, Lance Cabalona, Clayton Coker, Amber Glowacki, and Hannah Vahldick for help growing plants and making crosses. We would also like to thank Jeffrey Ross-Ibarra for help in calculating the divergence time between *ZmPME3* and *Pertunda*. The authors declare that they have no competing interests in this work. This work was supported by National Science Foundation Award number IOS-0951259 and by United States Department of Agriculture-National Research Initiative Competitive Grants Program Award number 35301-13314.

## Supplementary Materials

### Materials and Methods

#### Maize and teosinte lines and growth conditions

All maize and teosinte lines used in this study have been described previously (*3, 6, 7, 13*). Plants were grown under field conditions at either Stanford, California or Madison, Wisconsin.

#### *Tcb1-s* mapping

As described before (*3*), a Central Plateau teosinte collection 48703 (*31*) carrying the *Tcb1-s* barrier was backcrossed to the Mid-western US dent inbred W22 to incorporate the *Tcb1-s* locus into a maize background. This *Tcb1-s* strain was crossed to a chromosome 4 maize tester line *virescent17 (v17) brown midrib3 (bm3) sugary1 (su1)*, and the F1 was then backcrossed to the same tester line. Recombinants carrying crossovers between the four visual markers were tested for the *Tcb1-s* male and female functions in reciprocal crosses with *Tcb1-s/su1* F1 plants. PCR mapping markers were developed to refine the location of crossovers in these recombinants

#### *Tcb1-f* knockout mutant screen

To identify loss-of-function mutants of *Tcb1-female*, a *Ga1-m Tcb1-s* active *Mutator* strain was crossed to maize inbred A195 *su1 (tcb1)*, and then the progeny were grown as an open-pollinated block. Most of the progeny are expected to be heterozygous for *Tcb1-s* and *su1* in repulsion with *su1* approximately 6 cM away from the *tcb1* locus (*32*). Due to the rejection of the *Tcb1* pollen (which is predominantly su1), about 3% of the kernels in every ear with functional *Tcb1-f* are expected to be *sugary* in this open-pollinated population, while those without a crossing barrier were predicted to segregate *su1* at 25%.

#### Mixed pollination experiments

For the mixed pollination testing of the two *tcb1-f* knockout mutants, two pollen donor lines and three pollen receiver lines were used. Pollen from a maize line (*tcb1*) that does not have the *Tcb1* barrier genes but carries the endosperm color marker *R1-self color (R1-sc)* will produce purple kernel after fertilization of the lines used, while pollen from the knockout plants and the *tcb1-m* plants carry *r1-r* and produce anthocyaninless kernels that are white or yellow. After being collected from the two donors and mixed, pollen was put on the three receiving ears: (*1*) a *Tcb1-s* tester ear was used to verify the presence of the *tcb1* male function from the *tcb1-f::KO* pollen; (*2*) the *tcb1-f (KO)* ear was used to test the presence/absence of the female barrier function in the knockout mutant; and (*3*) a maize (*tcb1*) neutral ear was used to assay the percentage of viable pollen grains from the two donors in the mixture. The same protocol was used on the spontaneous *Tcb1-m* plant, except the *Tcb1-m* plant being tested was substituted for the *KO* plant. For mixed pollinations of the *Tcb1-f::silent lineage mop1* double mutant plants, pollen from the same *R1-sc tcb1* line and a *r1-r Tcb1-s* tester line was collected, mixed, and applied to the individual silent line ears and the neutral maize ears.

#### Silk tissue collection, RNA isolation and cDNA synthesis

Plants for RNA isolation were grown in summer field conditions in Stanford, CA. Silk tissues were collected around 11 am, immediately put into liquid nitrogen in the field, and stored at −80°C. Total RNA was isolated from silks with Trizol reagent (Invitrogen), DNase-treated, and either subjected to Illumina short read paired end RNA-seq, or used to synthesize the 1st strand cDNA with the Superscript IV RT kit (Invitrogen).

#### Quantitative RT-PCR

Each line/genotype had three biological replicates, and each in turn had three technical repeats. Tubulin (Zm00001d033850) was used as a reference gene. In each line, relative expression level of *Pertunda* was obtained by comparing *Pertunda* to tubulin.

#### Sequencing, assembly and analysis

All the RNA and DNA sequencing works were done with Illumina Paired-end sequencing by Novogene (CA, USA). RNA-seq reads from all samples were combined and *de novo* assembled with Trinity v2.4.0.(*33*) The gene in contig DN33598_c7_g3_i1 was identified as the *Tcb1-f* candidate gene due to its extremely high expression in the functional *Tcb1-s* line and the almost no expression in the *KO* mutants and a standard W22 maize line. PCR primers were designed based on the DN33598_c7_g3_i1 sequence, and one BAC clone was fished out from library made from maize line into which the *Tcb1-s::Col48703* haplotype had been introgressed. The BAC sequencing reads were assembled with SPAdes v3.11.1.(*34*) NODE_62, a contig that is 13656 bp with coverage of 4029, was identified as having the *Tcb1-f* candidate gene. Whole genome sequencing reads from the two *KO* mutants were individually assembled with SPAdes v3.11.1 and BLASTed against NODE_62. Also, the mutant sequencing reads were mapped against NODE_62 using GSNAP (*35*). Combining both approaches identified the *hopscotch* retrotransposon insertion in the *tcb1-f(KO1)* mutant allele.

For phylogenetic analyses, alignments were made using the ClustalW algorithm in MegAlign (DNASTAR). The predicted mature PME enzymes and the Arabidopsis PME family members were taken from Markovic and Janecek, as were the subfamily designations *(*26*)*. Phylogenies were produced from these alignments using MrBayes v3.2.0 using default settings for amino acid analysis (*36*). The MrBayes analysis was performed for 4,100,000 generations at which point the standard deviation of the split frequencies was below 0.004.

## Supplementary Figures

**Fig. S1.**
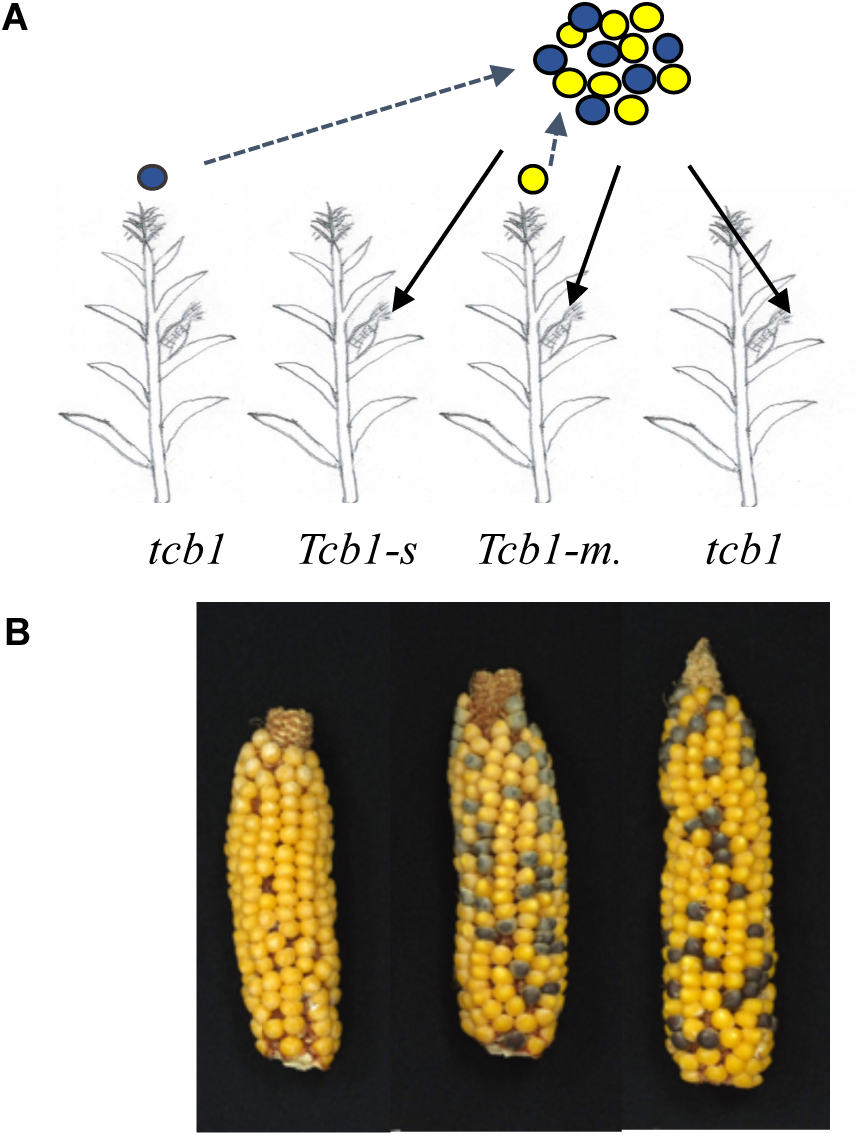
Mix pollination testing the spontaneous *Tcb1-m* plant. a. scheme of the experiment: Details in “Method” section. b. ears from the three pollen receivers for *Tcb1-m* plant test. Pollen from the *Tcb1-m* plant successfully fertilized the *Tcb1-s* ear and produced yellow kernels (left ear in b), while the ears from *Tcb1-m* plant indeed had lost the barrier to block maize pollen as shown by the purple kernels produced by *tcb1* pollen on the ears (middle ear in b).

**Fig. S2.**
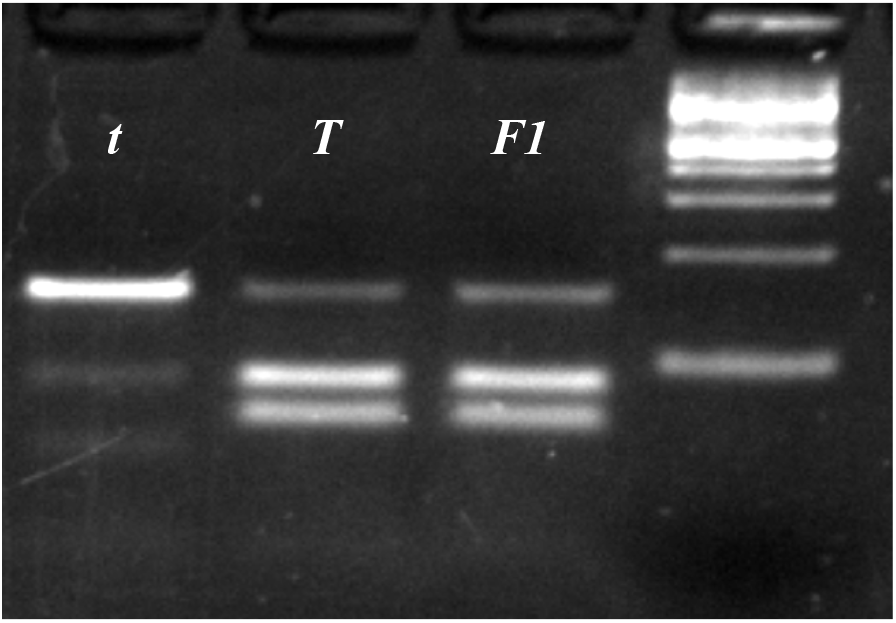
A dCAPS marker to test presence/absence of *Pertunda* in recombinants from the mapping population. The Marker was designed in the way that only the PCR amplicon from the *Tcb1* genomic DNA (*T*), but not the unspecific PCR product amplified from the maize gnomic DNA (*t*) would be cut by the enzyme Hae III. after PCR and enzyme digestion.

**Fig. S3.**
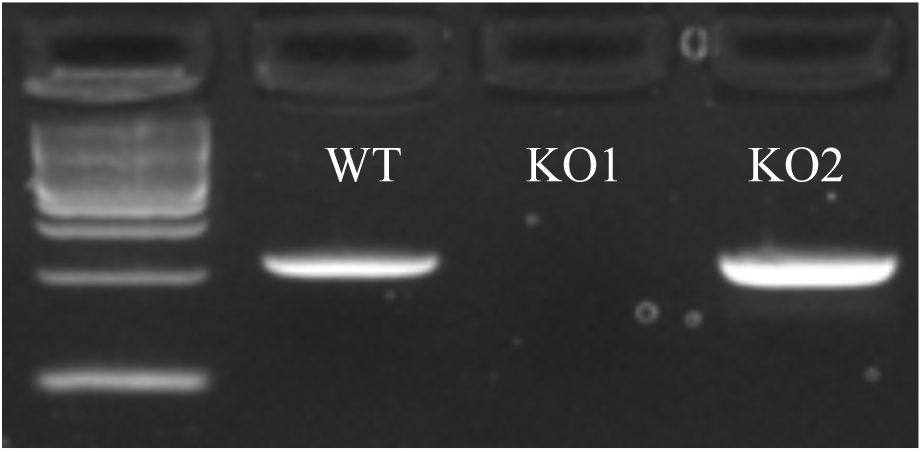
PCR detection of *Pertunda* in the two *tcb1-f(KO)* mutants. PCR primers spanning the possible mutation site in the *tcb1-f(KO1)* was designed and tested on the *Tcb1-s* (WT), *tcb1-f(KO1)* and *tcb1-f(KO2)*. Using *tcb1-f(KO1)* genomic DNA as template failed to produce amplicon, while *Pertunda* can be detected in the *tcb1-f(KO2)* mutant.

**Fig. S4.**
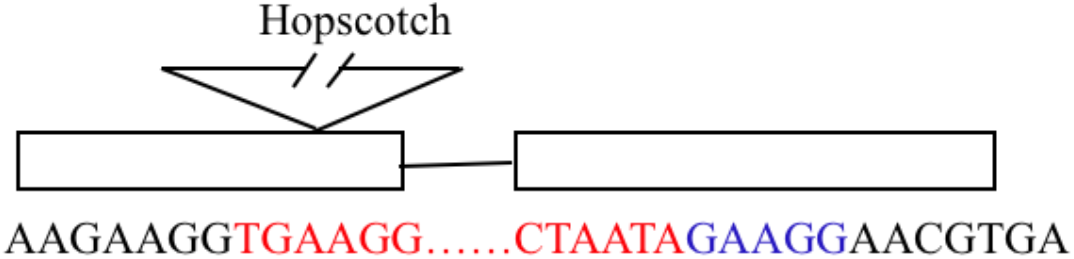
Identification of the mutation in the *tcb1-f(KO1)* mutant. Whole genome resequencing showed a Hopscotch retrotransposon insertion in the first exon. PCR primers based on the gene and the retrotransposon were used to confirm the insertion and the border sequences. Black bases, *Pertunda* gene sequence; Red bases, retrotransposon sequence; Blue bases, bases duplicated from the left border leading the retrotransposon sequence.

**Fig. S5.**
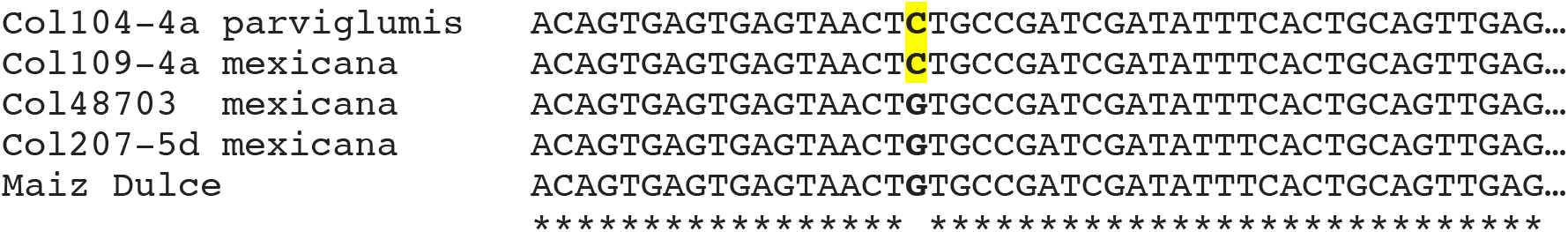
Partial DNA alignment of the *Pertunda* gene intron between the different *Tcb1-s* lines. Collections 109-4a, 48703 and 207-5d, ssp. *mexicana* teosintes; Collection104-4a, ssp. *parviglumis* teosinte; DGF1222, a traditional maize Dulce sweetcorn variety from Mexico (*13*).

**Fig. S6.**
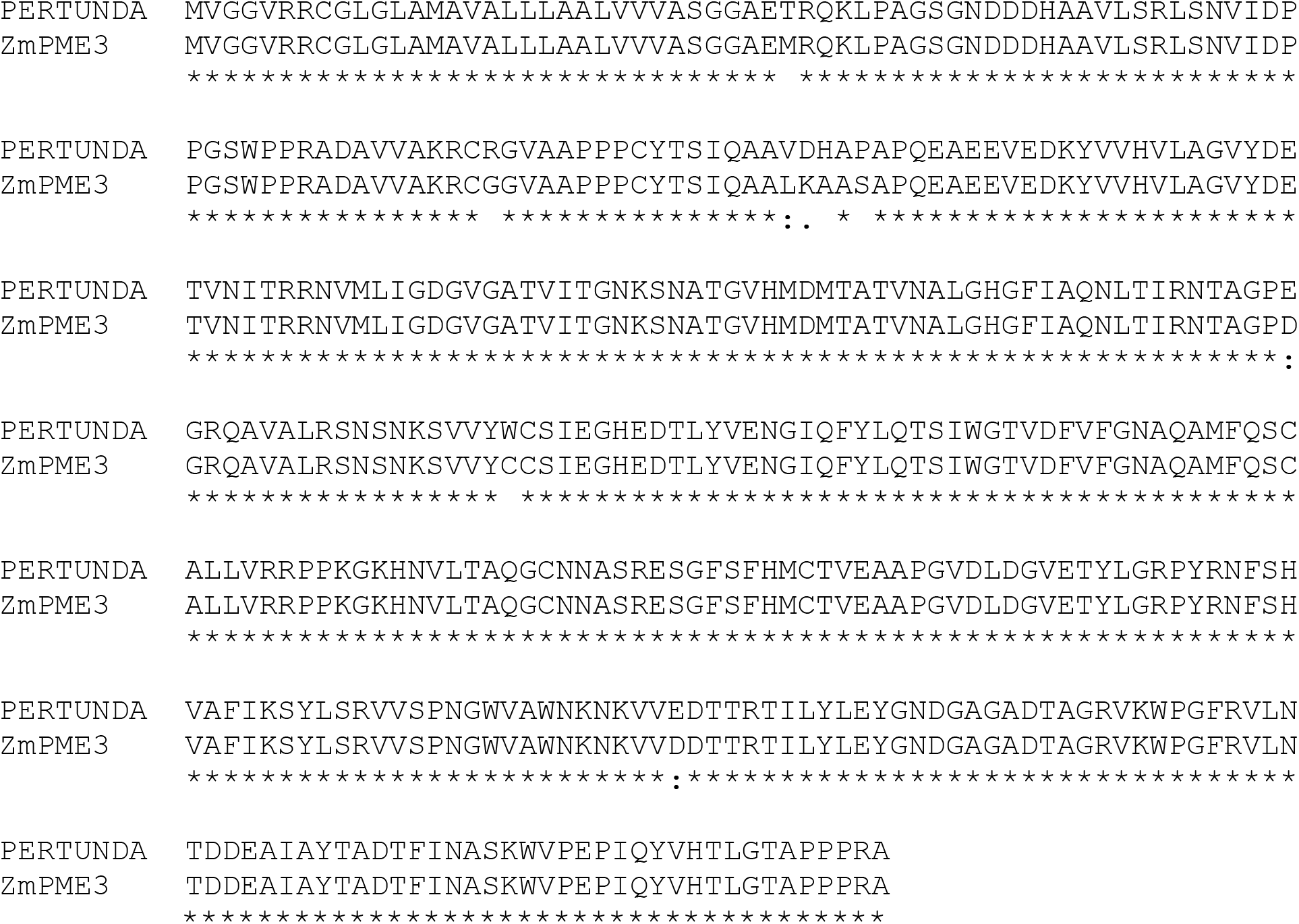
Alignment of Pertunda and ZmPME3 proteins with Clustal Omega.

**Fig. S7.**
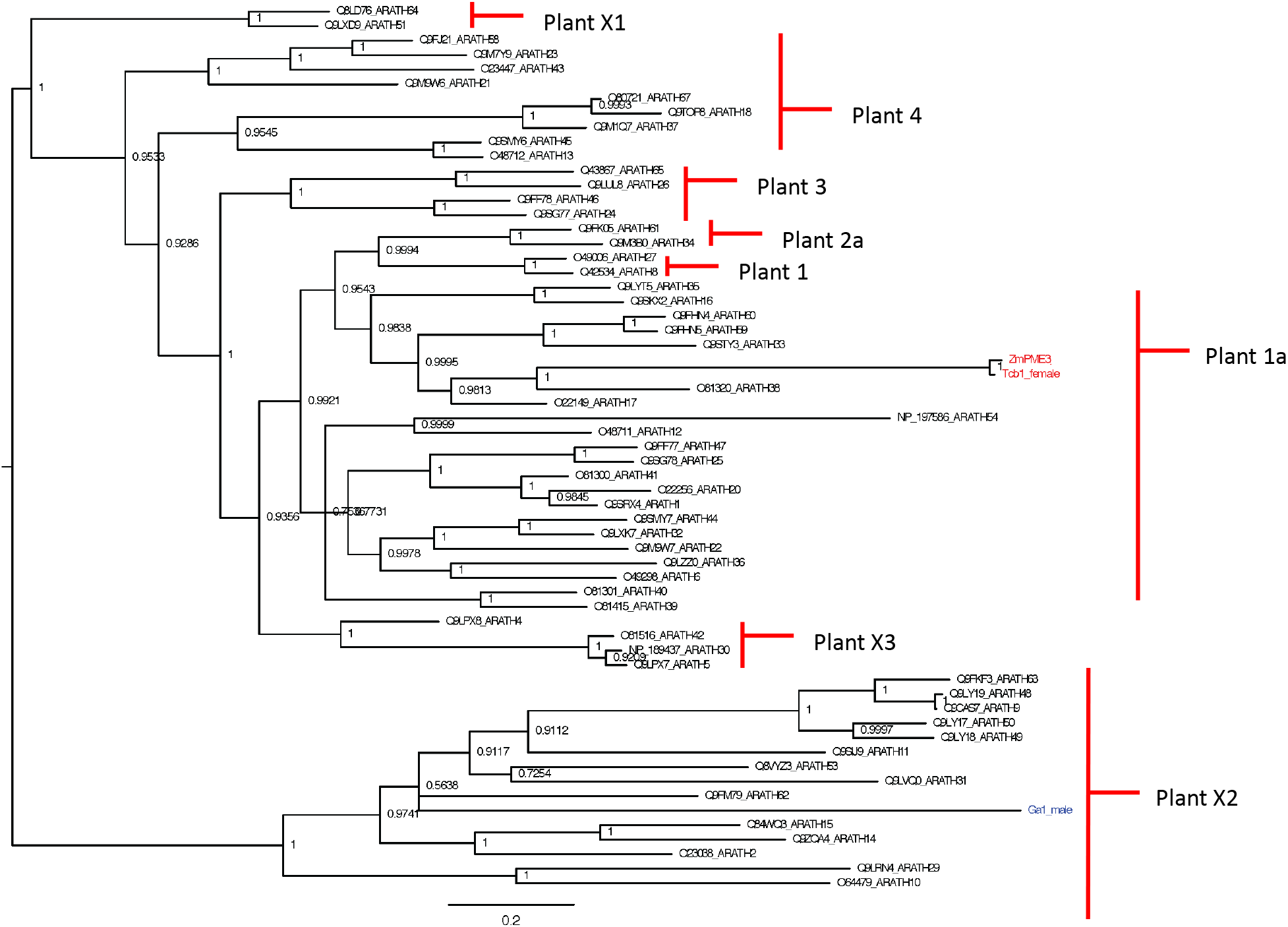
Phylogenic tree of mature PME enzymes (predicted pre and pro domains removed) of Arabidopsis PME proteins and predicted PME proteins encoded by cross-incompatibility loci of *Zea mays*.

**Table S1.**
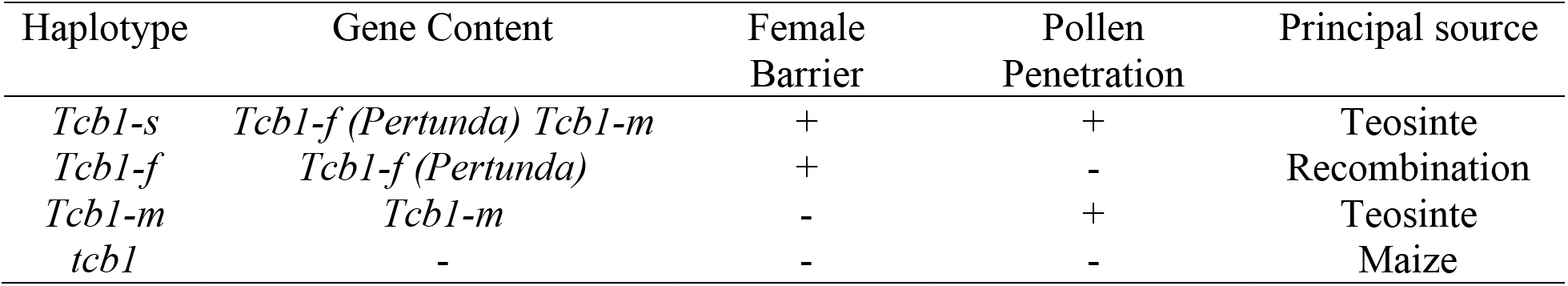
Haplotypes of the *tcb1* locus

**Table S2.**
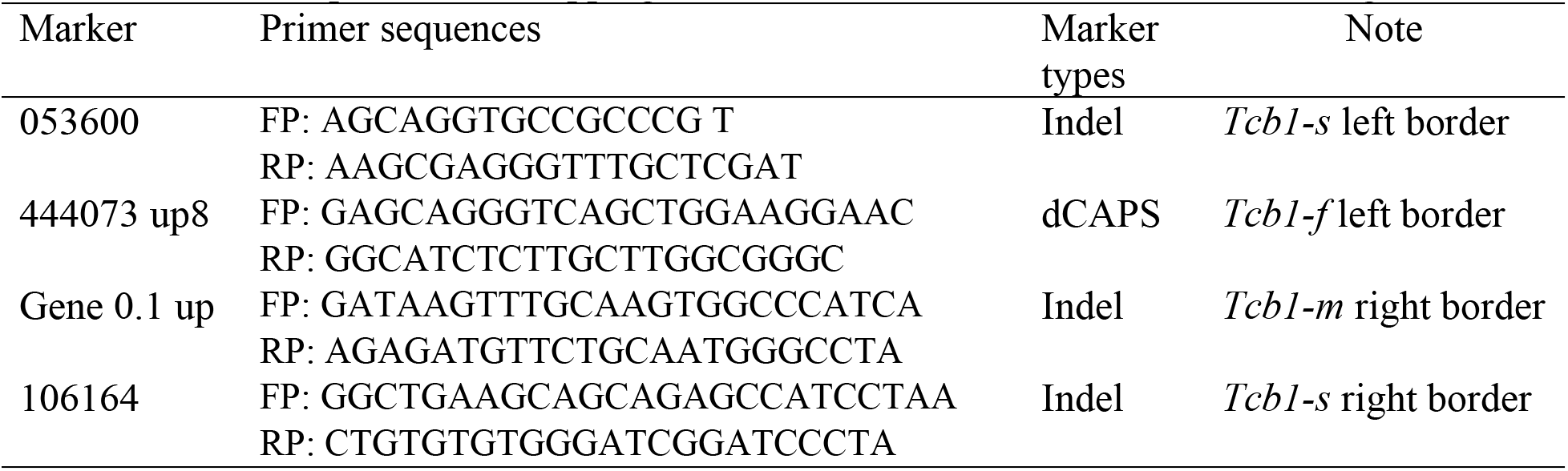
PCR primers for mapping *Tcb1-s* relative to the maize B73 reference genome

